# Preventing ovariectomy-induced weight gain decreases tumor burden in rodent models of obesity and postmenopausal breast cancer

**DOI:** 10.1101/2021.10.18.464856

**Authors:** Elizabeth A Wellberg, Karen A. Corleto, L. Allyson Checkley, Sonali Jindal, Ginger Johnson, Janine A. Higgins, Sarina Obeid, Steven M. Anderson, Ann D. Thor, Pepper J. Schedin, Paul S. MacLean, Erin D. Giles

## Abstract

Obesity and adult weight gain are linked to increased breast cancer risk and poorer clinical outcomes in postmenopausal women, particularly for hormone-dependent tumors. Menopause is a time when significant weight gain occurs in many women, and clinical and preclinical studies have identified menopause (or ovariectomy) as a period of vulnerability for breast cancer development and promotion. We hypothesized that preventing weight gain after ovariectomy (OVX) may be sufficient to prevent the formation of new tumors and decrease growth of existing mammary tumors. Here, we tested this hypothesis in a rat model of obesity and carcinogen-induced postmenopausal mammary cancer and validated our findings in a murine xenograft model with implanted human tumors. In both models, preventing weight gain after OVX significantly decreased obesity-associated tumor development and growth. Importantly, we did not induce weight loss in these animals, but simply prevented weight gain. In both lean and obese rats, preventing weight gain reduced visceral fat accumulation and associated insulin resistance. Similarly, the intervention decreased circulating tumor-promoting growth factors and inflammatory cytokines (ie, BNDF, TNFα, FGF2), with greater effects in obese compared to lean rats. In obese rats, preventing weight gain decreased adipocyte size, adipose tissue macrophage infiltration, reduced expression of the tumor-promoting growth factor FGF-1, and reduced phosphorylated FGFR in tumors. Together, these findings suggest that the underlying mechanisms associated with the anti-tumor effects of weight maintenance are multifactorial, and that weight maintenance during the peri-/post-menopausal period may be a viable strategy for reducing obesity-associated breast cancer risk and progression in women.

## INTRODUCTION

Obesity is a global epidemic, affecting >640 million adults worldwide and rates continue to rise (1–3). Obesity rates are higher for women than men, and obesity is especially prevalent in women in the peri-menopausal or early postmenopausal life stages (45-65 years) compared to younger women. Excess adiposity increases both breast cancer risk and cancer-specific mortality, and these effects are modulated by menopausal status (4–9). Prior to menopause, obesity’s effects are generally considered negligible or even protective, particularly for estrogen-receptor positive (ER+) tumors (10,11). However, after menopause obesity increases the incidence, progression, and eventual mortality from breast cancer by up to 40% compared to women at a healthy weight (12). The risk is notably elevated in women with a history of weight gain throughout life, as well as those who gain weight in the 5-6 year period immediately preceding their breast cancer diagnosis (13–15). Given the prevalence of obesity and breast cancer, it is critical to determine effective interventions that can prevent tumor development and disease progression in the context of excess adiposity.

To study the link between obesity and postmenopausal ER+ tumor growth, we developed the OR/OP-OVX model (16–20) which is highly reflective of many aspects of obesity-associated postmenopausal breast cancer. Specifically, the adverse impact of obesity emerges during the brief period of rapid weight gain induced by ovariectomy (OVX). Prior to menopause, obesity is associated with metabolic inflexibility or insulin resistance. While insulin resistance has been identified as a likely mechanism linking obesity and tumor promotion (21–26), the independent impact of insulin resistance *prior to menopause* is relatively modest suggesting that this remains relatively inert until menopause. We have proposed a dual-requirement hypothesis, whereby the combination of obesity-associated metabolic dysfunction and menopause-induced weight gain together creates an environment conducive to tumor development and growth. In support of this hypothesis, our previous studies found that treating rats with the antidiabetic drug metformin during the post-OVX period improved underlying metabolic dysfunction (20) and significantly reduced both the growth of existing tumors and the development of new tumors, without impacting weight gain (17,20). The goal of the current study was to determine if preventing weight gain during the post-OVX period was sufficient to decrease obesity-associated tumor development and growth in a setting that models menopause.

It has been clearly established that diet-induced weight loss or weight loss through bariatric surgery both decrease breast cancer risk and improve outcomes in patients with breast cancer. However, there is overwhelming evidence that weight loss is difficult to sustain (27), and menopause is a time when women are particularly prone to gaining weight. Therefore, our goal in the current study was not to induce weight loss in animals, but instead to maintain animals at their pre-OVX weight and prevent the weight gain induced by OVX as a model for preventing menopausal weight gain (referred to as weight maintenance). Here, we show that this weight-maintenance approach significantly decreased mammary tumor burden in both obese and lean animals, and also prevented the formation of new tumors after OVX. These beneficial effects of weight gain prevention had positive impacts on systemic metabolic and inflammatory markers, the tumor microenvironment, and directly on tumors, which resulted in a lower incidence and improved outcomes. These findings suggest that the peri-menopausal/menopausal period of weight gain may provide an ideal “window of opportunity” for interventions aimed at improving cancer outcomes (16,17). Given this finding, clinical studies that focus on prevention of weight gain may be highly beneficial, even without weight loss, in women at risk for postmenopausal breast cancers.

## METHODS

All animals used in these studies were housed at 22–24°C with a 12:12-h light-dark cycle and free access to water. All procedures were approved by the appropriate Institutional Animal Care and Use Committee.

### Rat Model of Obesity and Breast Cancer

Our OP-OR/OVX model of obesity and postmenopausal breast cancer was used, as previously described (18,28). We, and others, have shown that tumors that develop using this method are similar to human breast tumors with regard to: (a) the percentage of tumors that are intraductal, (b) the progression of histologic stages from hyperplasia, to carcinoma in situ, to invasive cancer, and (c) steroid receptor status (16,17,29).

Female Wistar rats (100–125 g; 5 weeks of age) were purchased from Charles River Laboratories (Wilmington, MA). Rats were individually housed in wire bottom cages to limit physical activity and were given ad libitum access to purified high-fat diet (HF; 46% kcal fat; Research Diets #D12344, New Brunswick, NJ) to induce obesity in this genetically susceptible strain. All rats remained on the HF diet for the duration of the study. Animals were ranked by their percent body fat at the time of OVX surgery (mean 27.1 weeks of age). Rats in the top and bottom tertiles of adiposity were classified as obese and lean, respectively. Rats from the middle tertile were removed from this study.

To induced mammary tumor formation, 55-day old female rats (+/- 1d) were given a single intraperitoneal injection of the carcinogen N-methylnitrosourea 1-methyl-1-nitrosourea (MNU, 50 mg/kg; #MRI-340, MRI Global, Kansas City, MO) Tumors were monitored by manual palpation and measured weekly with digital calipers for the duration of the study. Tumor volumes were calculated as π * (length/2 * width/2 * height/2).

Body weight and food intake were monitored weekly, as previously described (30,31).

Body composition was determined at 18 weeks of age, on the day of OVX, every 2 weeks post-OVX, and again at the time of sacrifice, by quantitative magnetic resonance (qMR; EchoMRI; Echo Medical Systems, Houston, TX). In a rolling study design, rats underwent surgical ovariectomy (OVX) to mimic the post-menopausal state once they developed at least one mammary tumor ≥ 1cm^3^. OVX surgery was performed under isoflurane anesthesia. At the time of OVX, animals were randomly assigned to either be maintained at their pre-OVX body weight (weight-maintained; WM n=10 lean and 15 obese) or ad libitum fed (AdLib; n= 23 lean and 17 obese) for the remainder of the study. WM rats were maintained at their pre-OVX body weight by providing a limited portion of the HF diet each day, immediately prior to the start of the dark cycle. Each rat was weighed daily and adjustments to the food allotment were made if a 2-3 day trend of weight loss or gain occurred. Rats were euthanized by exsanguination under anesthesia 8 weeks after OVX, or when tumor burden exceeded 10% of the animal’s body weight.

### Plasma Measurements

Tail vein blood was collected at the time of OVX, at 2 weeks post-OVX, and again at the time of sacrifice. Blood was drawn during the latter part of the light cycle; plasma was isolated and stored at −80°C. Plasma insulin was measured by ELISA (Alpco 80-INSRT-E01, Salem, NH). Colorimetric assays were used to measure plasma free fatty acids (Wako Chemicals USA, Richmond, VA), glucose, triglycerides (TG), and total cholesterol (#TR15421, TR22321, and TR13521, respectively; Thermo Fisher Scientific, Waltham, MA). Inflammatory markers were measured using a 90-plex antibody array (Rat L90 Array, AAR-SERV-LG, RayBiotech Life, Inc., Peachtree Corners, GA).

### Histological staining and imaging

Sections of formalin fixed paraffin embedded tissue (4 μm) were stained with hematoxylin and eosin (H&E) using a Sakura autostainer (Sakura Finetek, Torrance, CA, USA). Mammary tumors were classified histologically by the criteria of Young and Hallowes (32) and only adenocarcinomas were included in subsequent analyses. For immunohistochemical detection of proteins, tissue sections were stained with antibodies targeting adipophilin (LS-C348703, Lifespan Biosciences) at 1:300 dilution for 60 minutes followed by mouse on rat secondary antibody (MRT621H, Biocare) for 30 min and DAB chromogen or phospho-FGFR1 (Y654; Abcam ab59194 1:500) followed by ImmPRESS^®^ HRP Horse Anti-Rabbit IgG Polymer Detection Kit (Vector Laboratories, MP-7401). All slides were counterstained with hematoxylin (S330130, Dako, Carpinteria, CA).

Stained slides were scanned using an Aperio slide scanner (Leica Biosystems, Buffalo Grove, IL) at 20X magnification, corresponding to 0.43 μm per pixel which enables high resolution access to the entire tissue section via a virtual image. Images were evaluated using Imagescope software and signal captured. Liver adipophilin was quantified using Aperio algorithms. pFGFR1 was manually scored in blinded samples as a percentage of positive cells in 10% increments.

### Adipocyte Cellularity

H&E-stained slides were used to assess adipocyte cell size distribution and mean adipocyte diameter. Using Imagescope software, 5 regions of each tissue section were randomly selected for analysis. Images were exported, and cell diameter and number were determined using the Adiposoft plug-in for ImageJ (FIJI).

### Tissue Analysis (PCR)

Total RNA was isolated from pulverized mammary adipose or tumor tissues using TRIzol reagent (Thermo Fisher Scientific) according to manufacturer’s protocol. Q-RT-PCR was performed using TaqMan primers/probe sets (Applied Biosystems) and analyzed as transcript copies per 50 ng RNA expressed relative to RNAPII expression as previously described (33).

### Mouse Model of Obesity and Human Xenograft Tumors

Female Rag1-null (Jackson Labs Stock #002216) were fed a high fat high sucrose diet (45%; Research Diets #D15031601) for approximately 16 weeks (28). They were then ovariectomized and supplemented with 0.5 uM 17ß-estradiol (E2) in the drinking water (28). A 2 mm x 3 mm fragment of the ER-positive patient derived breast tumor, UCD12 (34), was implanted into the inguinal mammary fat pads and allowed to reach approximately 0.5 cm in diameter(33). At this time, mice were randomized based on body fat percentage to AdLib or WM intervention groups as described for the rat study above. Supplemental E2 was withdrawn, and body weight was recorded daily. WM mice were fed only enough of the HF diet to prevent weight gain. The study ended and tissues were harvested after 18 days of treatment.

### Statistical analysis

Data were analyzed with SPSS 26.0 software or using GraphPad Prism v9. Where applicable data are expressed as mean ± standard error of the mean (SEM). Comparisons between two groups were assessed by t tests. When comparing more than two groups, P values were assessed using two-way ANOVA, examining the effect of adiposity (lean vs obese), weight-maintenance intervention (weight-maintained vs control), and the interaction of the two. In some cases, data were analyzed by analysis of covariance with a specified covariate in the model. Relationships between variables were assessed with the Spearman correlation coefficient.

## RESULTS

### Baseline Rat Characteristics

Our previous work identified both OVX-induced overfeeding (positive energy imbalance) and obesity-associated metabolic dysfunction (peripheral insulin resistance) as important potential drivers of mammary tumor progression (17). Based on this finding, the goal of the current study was to determine if preventing OVX-induced weight gain without promoting weight loss, would be sufficient to inhibit tumor progression and development of new tumors. An overview of the study design is shown in **Fig 1A**.

**Figure 1.**
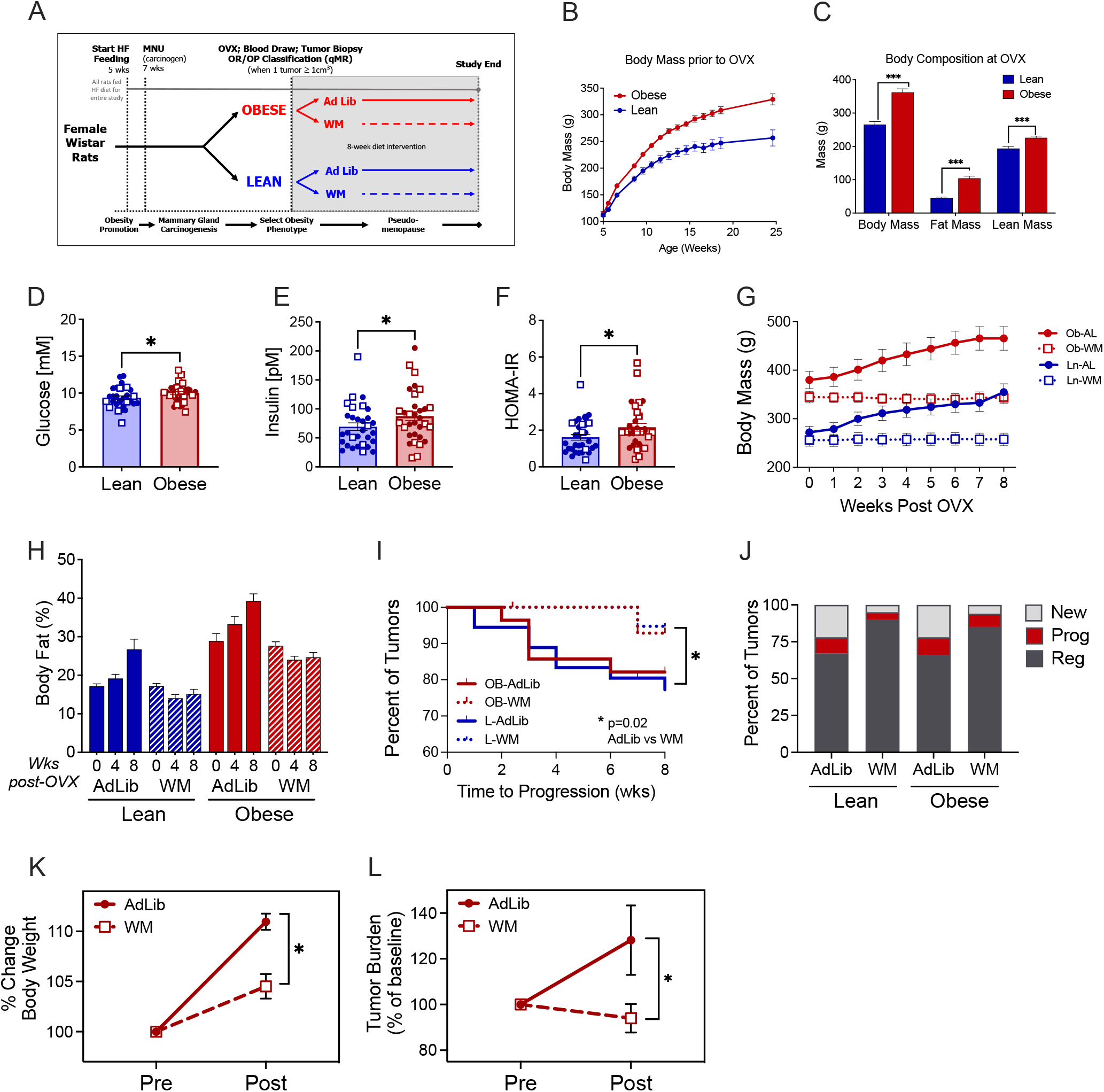
Preventing OVX-induced weight gain improves mammary tumor outcomes in lean and obese rats. **A)** Study design diagram depicting times at which rats were given high fat diet (HFD), N-methyl, N-nitrosourea (MNU), and randomized to *ad libitum* (Ad Lib) feeding or weight maintenance (WM) intervention. **B)** Body mass of lean and obese rats prior to OVX. Data are mean ± SEM. **C)** Body composition of lean and obese rats at OVX, measured by qMR. Unpaired t-test, P<0.001. **D-F)** Blood was collected from fasted rats at OVX and glucose (D) and insulin (E) were measured and used to calculate HOMA-IR (F). Filled circles (●) indicate rats assigned to AdLib, open squares (■) indicate rats assigned to WM. Unpaired t-test, P<0.05. **G)** Body mass of lean and obese AdLib (AL ●) or WM (■) rats beginning at OVX and continuing for the 8-week intervention. **H)** Body fat percent of lean and obese AdLib and WM rats measured by qMR at 0-, 4-, and 8-weeks post OVX. **I)** Kaplan-Meier survival curve showing the time to progression of existing tumors at OVX in lean and obese AdLib or WM rats. Log rank P=0.02. **J)** Percent of tumors that were existing at OVX and progressed (red) or regressed (dark grey), or that were new after OVX (light grey). **K-L)** (K) Body mass and (L) tumor burden of UCD12 patient-derived xenograft tumors in OVX AdLib or WM Rag1-null mice fed a high fat/high sucrose diet measured before and 3 weeks after estrogen withdrawal.

Lean and obese phenotypes were defined based on percent body fat at the time of OVX. When analyzed retrospectively, obese rats had significantly higher body weight starting as early as 6 weeks of age, compared to lean rats (**Fig 1B**). Similarly, percent body fat was significantly higher in the obese as early as 9 weeks and remained elevated at 14 weeks (**Supplementary Table 1**). At the time of OVX surgery, the 29% greater body weight in obese rats was primarily due to a doubling of fat mass in the obese, with only a small increase in lean mass (**Fig 1C**). When expressed as a percentage of body weight, body fat remained higher in the obese when compared to their lean counterparts (27.6 ± 0.9% vs 17.3 ± 0.4%), replicating our previous studies using this model (16–20,31).

Fasting glucose and insulin were measured at the time of OVX, and the homeostatic model assessment of insulin resistance (HOMA-IR) was calculated. Obese rats had significantly higher fasting glucose (**Fig 1D**) and insulin (**Fig 1E**), which resulted in significantly elevated HOMA-IR (**Fig 1F**) when compared to lean rats. There were no differences between those randomized to the WM or AdLib groups (**Fig 1D-F**, circles vs squares).

### Preventing OVX-Induced Weight Gain in Lean and Obese Rats

As expected after the loss of ovarian hormones following OVX (16–18,28), ad libitum fed control rats gained weight rapidly over the early post-OVX period independent of pre-OVX obesity status, with a cumulative 50.5 ± 3.2g gained in the first 4 weeks after OVX, with no difference between lean and obese animals (**Fig 1G** and **Supplementary Table 2**). All AdLib fed rats continued to gain weight and increase in adiposity across the 8-week post-OVX period. At the end of the study, obese rats remained heavier than their lean counterparts (438.2 vs 352.3g), with greater fat and lean mass (**Fig 1H**). Despite the higher absolute body weight in the obese rats, the lean and obese gained similar amounts of weight and body fat when expressed relative to their pre-OVX levels. In the 8 weeks after OVX, body weight increased by 27 and 24% in lean and obese AdLib rats, respectively (**Fig 1H**). Importantly, across the 8-week diet intervention, WM rats were maintained within 3% of their OVX weight (**Fig 1G**), with no significant change in body composition after OVX (**Fig 1H**).

### Preventing Weight Gain after OVX Improves Tumor Outcomes

Lean and obese rats entered the OVX phase of the study with no difference in the number of tumors per animal (mean = 1.83 ± 0.13) or total tumor burden (mean = 2.33 ± 0.24). Both lean and obese WM rats had a significantly longer time to tumor progression compared to AdLib controls (**Fig 1I**), with no differences between adiposity groups. At the end of the 8-week intervention, tumors were classified as existing at OVX and progressing (growing), existing at OVX and regressing (shrinking or unchanged), or newly formed after OVX. In both lean and obese groups, the WM rats had fewer tumors that progressed, more tumors that regressed, and developed fewer new tumors compared to AdLib rats (**Fig 1J**). We found similar beneficial anti-tumor effects of preventing OVX-induced weight gain in a confirmatory mouse xenograft model (**Fig 1K**). Together, these data indicate that OVX-induced weight gain is tumor promotional regardless of adiposity status (lean vs obese) and preventing weight gain during this window of time prevents the growth and progression of tumors, even if obesity is not reversed.

### Preventing Weight Gain Improves Systemic Metabolic Markers

There are several mechanisms by which preventing weight gain or preventing adipose tissue expansion may improve tumor outcomes, including systemic effects, and direct effects on the tumor and/or the tumor microenvironment. We evaluated potential systemic effects by measuring plasma metabolites, inflammatory markers, and both hepatic and adipose makers of metabolic health. Importantly, all animals in this study consumed a HF diet for the duration of the study, which was reflected in fasting metabolite levels that were higher than what is typically see in rodents on a low fat or chow diet.

#### Plasma Metabolites

Glucose, insulin, and HOMA-IR were assessed at the end of the 8-week intervention as markers of whole-body insulin resistance. Fasting glucose and insulin were lower in the WM groups compared to controls, regardless of adiposity (**Supplemental Fig 2**). When glucose and insulin were used to calculate HOMA-IR, this measure of insulin resistance was also significantly lower in WM rats compared to AdLib controls (**Supplemental Fig 2**). Repeated measures analysis demonstrated a significant time by treatment effect, demonstrating increased glucose, insulin, and HOMA-IR in the AdLib groups from pre to post-OVX and reduction in the WM groups across this same time (**Figs 2A-C**). Similarly, obese animals also had higher circulating, cholesterol, triglycerides (TG), and a trend for higher non-esterified fatty acids (p = 0.10) than their lean counterparts; however, only circulating cholesterol was significantly reduced in the WM animals (**Supplemental Table 3**). Reflective of adiposity levels, circulating leptin was also significantly higher in obese relative to lean group rats and was further reduced in the WM groups (**Supplemental Table 3**).

**Figure 2.**
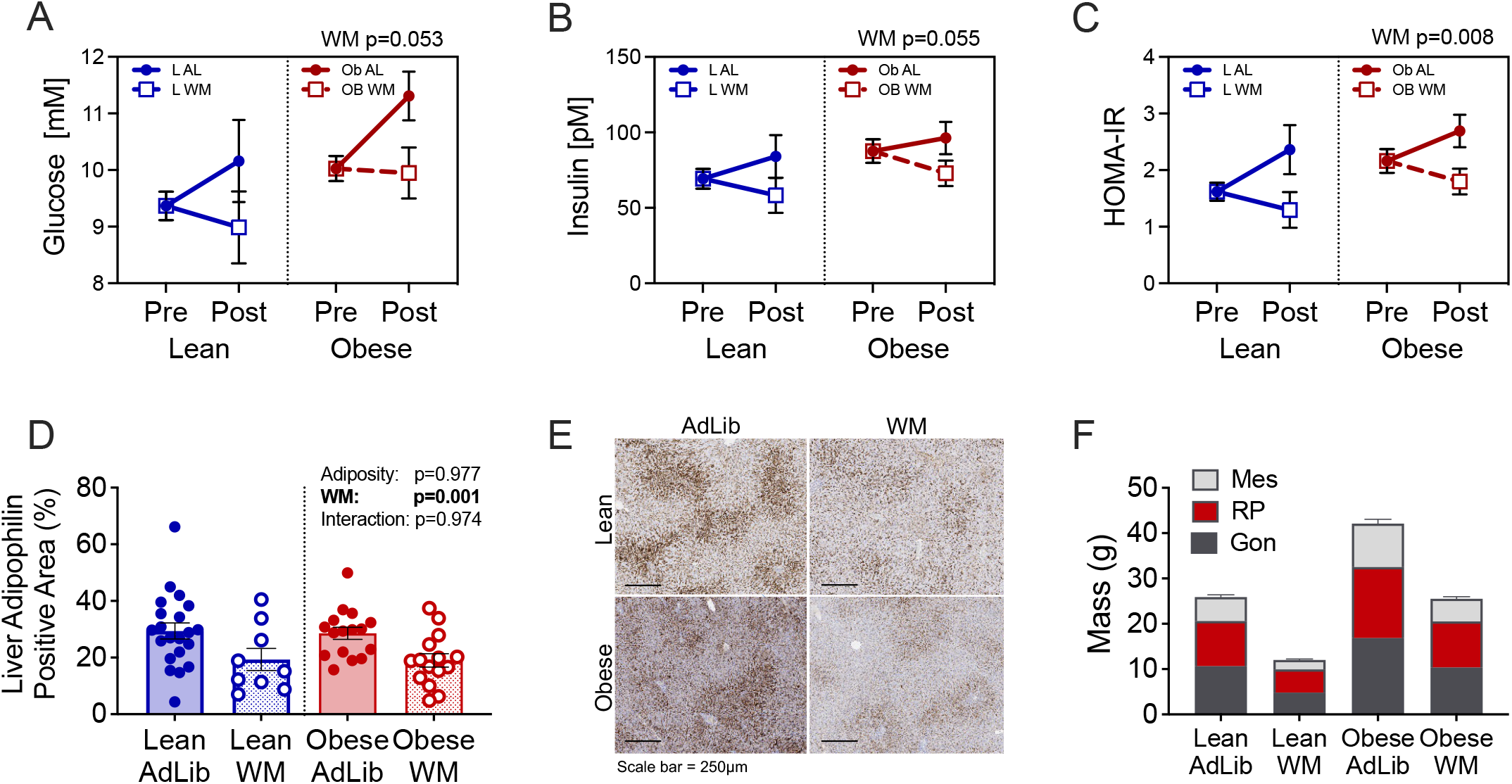
Weight Maintenance after OVX improves markers of metabolic function in lean and obese rats. **A-C)** Blood was collected from fasted rats at the end of study and glucose (A) and insulin (B) were measured and used to calculate HOMA-IR (C). Filled circles indicate rats assigned to AdLib, open squares indicate rats assigned to WM. Two-way ANOVA, main effects of adiposity or WM. **D)** Quantification of adipophilin in liver sections from lean and obese AdLib or WM rats at the end of study, measured by IHC. Two-way ANOVA, main effects of adiposity or WM. **E)** Representative images of liver adipophilin staining. Scale bar is 250 μm. **F)** Mass of visceral [mesenteric (Mes; light grey), retroperitoneal (RP; red), and gonadal (Gon; dark grey)] adipose depots in lean and obese AdLib or WM rats at the end of study.

#### Hepatic Lipid Accumulation and Fat Distribution

Hepatic lipid accumulation is indicative of impaired metabolic health; therefore, we assessed the impact of WM on hepatic steatosis. Hepatic lipids were measured using semi-quantitative IHC analysis of adipophilin (a marker of lipid droplet membranes). We have previously shown that both lean and obese rats have low levels of hepatic lipid prior to OVX, which increases significantly following OVX-induced weight gain in all animals (20). Here, adipophilin staining was significantly reduced in WM rats compared to AdLib rats, regardless of adiposity (**Fig 2D**), further supporting the beneficial effects of WM on metabolic health.

Increased visceral fat accumulation, which occurs during menopause (35), has also been associated with several features of metabolic disease, including insulin resistance and systemic inflammation (36). Thus, in addition to measuring total body fat, we also assessed regional fat distribution by weighing fat pads at the end of the study. As expected, AdLib rats had significantly larger visceral fat depots (gonadal, retroperitoneal, and mesenteric fat pads; **Fig 2E**) than their WM counterparts, regardless of whether they were lean or obese. The WM intervention resulted in significant fat loss in each depot, suggesting that this intervention reduced visceral fat gain globally. Together, the beneficial effects of WM on glucose, insulin, hepatic steatosis, and visceral adiposity indicates that preventing weight gain during this relatively short post-OVX period improves metabolic health, even without reversal of obesity.

#### Plasma Inflammatory Cytokines & Growth Factors

To better understand the beneficial effects of WM on whole-body and tumor outcomes, we performed targeted proteomics analysis of plasma collected at the end of the 8-week intervention period from AdLib and WM groups. Of 90 total proteins analyzed, 24 were modulated similarly by WM in both lean and obese rats (p<0.1; **Supplemental Table 4**). We separated these into those increased with WM compared to those that were decreased. Among the top KEGG pathways significantly increased in WM rats were several inflammatory pathways (cytokine/cytokine receptor interactions, IL-17 signaling, asthma, intestinal immune network, TNF signaling, and chemokine signaling), as well as pathways known to play a role in tumor growth (Jak/Stat signaling and pathways in cancer) (**Fig 3A**). Examples of proteins that contributed to these pathways included: interleukin-2 (IL-2), macrophage inflammatory protein 2 (MIP2/CXCL2), and macrophage inflammatory protein 1α (MIP-1α/Ccl3) (**Fig 3B-D**). Pathways that were decreased in the WM group included lipid-related pathways (adipocytokine signaling, non-alcoholic fatty liver disease), as well as pathways known to support tumor growth (MAPK, PI3K/Akt, Ras, JAK/STAT signaling) (**Fig 3E).** Examples of proteins that contributed to these pathways included: brain-derived neurotrophic factor (BDNF), tumor necrosis factor alpha (TNF-α), and fibroblast growth factor 2 (FGF2) (**Fig 3 F-H**). These data suggest that beneficial effects of the weight-maintenance intervention are likely due, in part, to the changes in these systemic inflammatory cytokines and growth factors.

**Figure 3.**
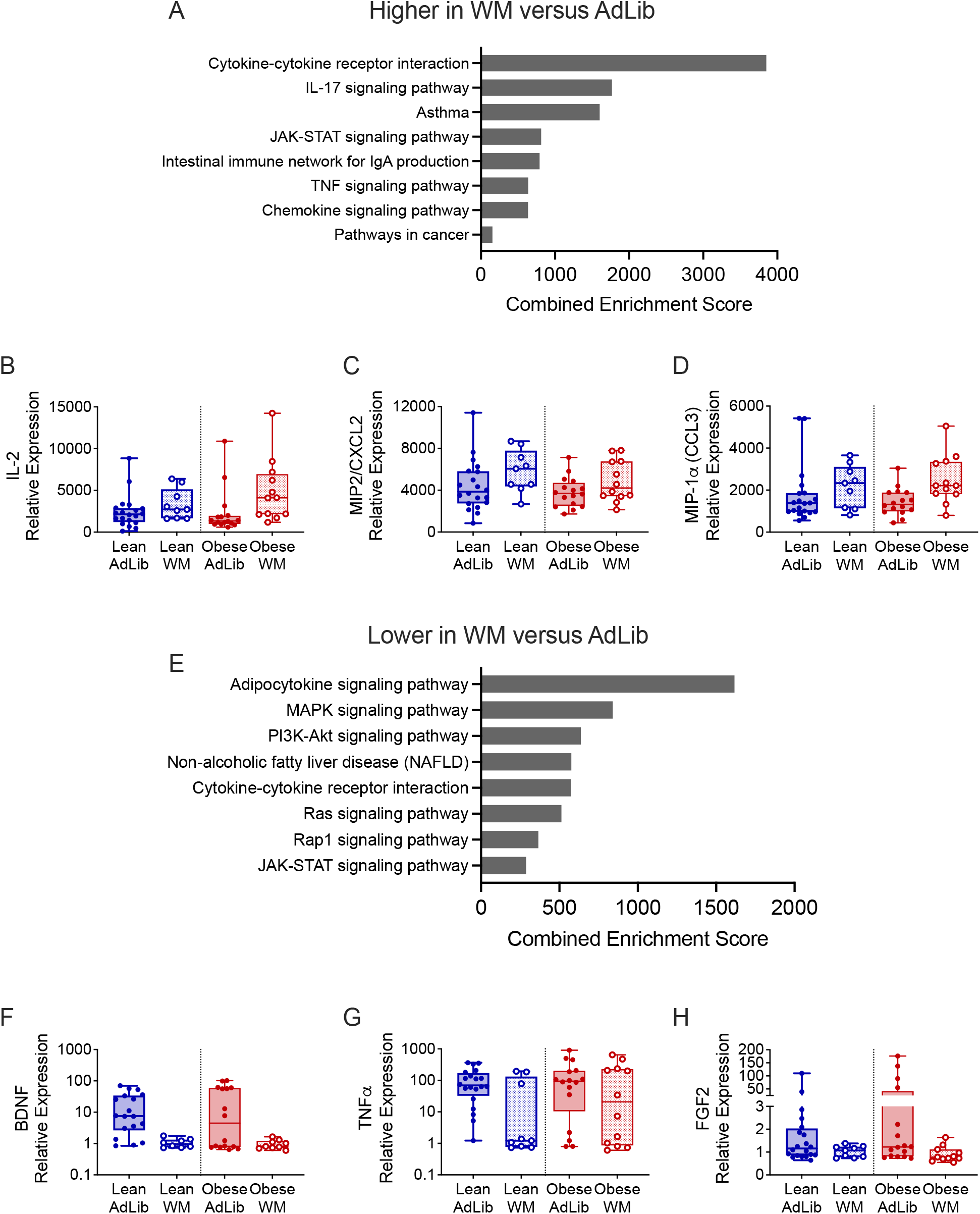
Weight Maintenance after OVX alters plasma proteins associated with tumor progression and metabolism in lean and obese rats. **A)** GO Biological Processes common to plasma cytokines that were higher (p<0.1) in WM versus AdLib lean and obese rats at the end of study. **B-D)** Examples of cytokines that were greater in plasma from WM versus AdLib rats, including IL-2 (B), MIP2/CXCL2 (C), and MIP-1α /CCL3 (D). **E)** GO Biological Processes common to plasma cytokines that were lower (p<0.1) in WM versus AdLib lean and obese rats at the end of study. **F-H)** Examples of cytokines or growth factors that were lower in plasma from WM versus AdLib rats, including BDNF (F), TNFα (G), and FGF2 (H).

### Preventing Weight Gain Decreases Tumor Promoting Potential of the Local Mammary Adipose Microenvironment

Expansion of adipose depots during weight gain can involve adipocyte hypertrophy and/or hyperplasia. Adipocyte hypertrophy has been directly linked with the development of insulin resistance and growth factor signaling (37). Based on the decrease in both insulin resistance and circulating inflammatory proteins and growth factors by WM in lean and obese rats, we evaluated adipocyte size distribution in mammary (subcutaneous) adipose tissue, which is a component of the tumor microenvironment. As shown in **Fig 4A,** Obese AdLib rats had fewer small adipocytes (20-60 μm in diameter) and more large adipocytes (60-120 μm) than lean AdLib rats (**Fig 4A**). As expected, the WM intervention reduced the proportion of large adipocytes and increased the proportion of small adipocytes in both the lean and obese rats (**Fig 4A,** dotted lines) compared to AdLib controls (**Fig 4A**, solid lines). This resulted in a significantly lower mean adipocyte diameter in subcutaneous adipose in the obese WM vs AdLib group, with a similar trend in the lean (**Fig 4B**).

**Figure 4.**
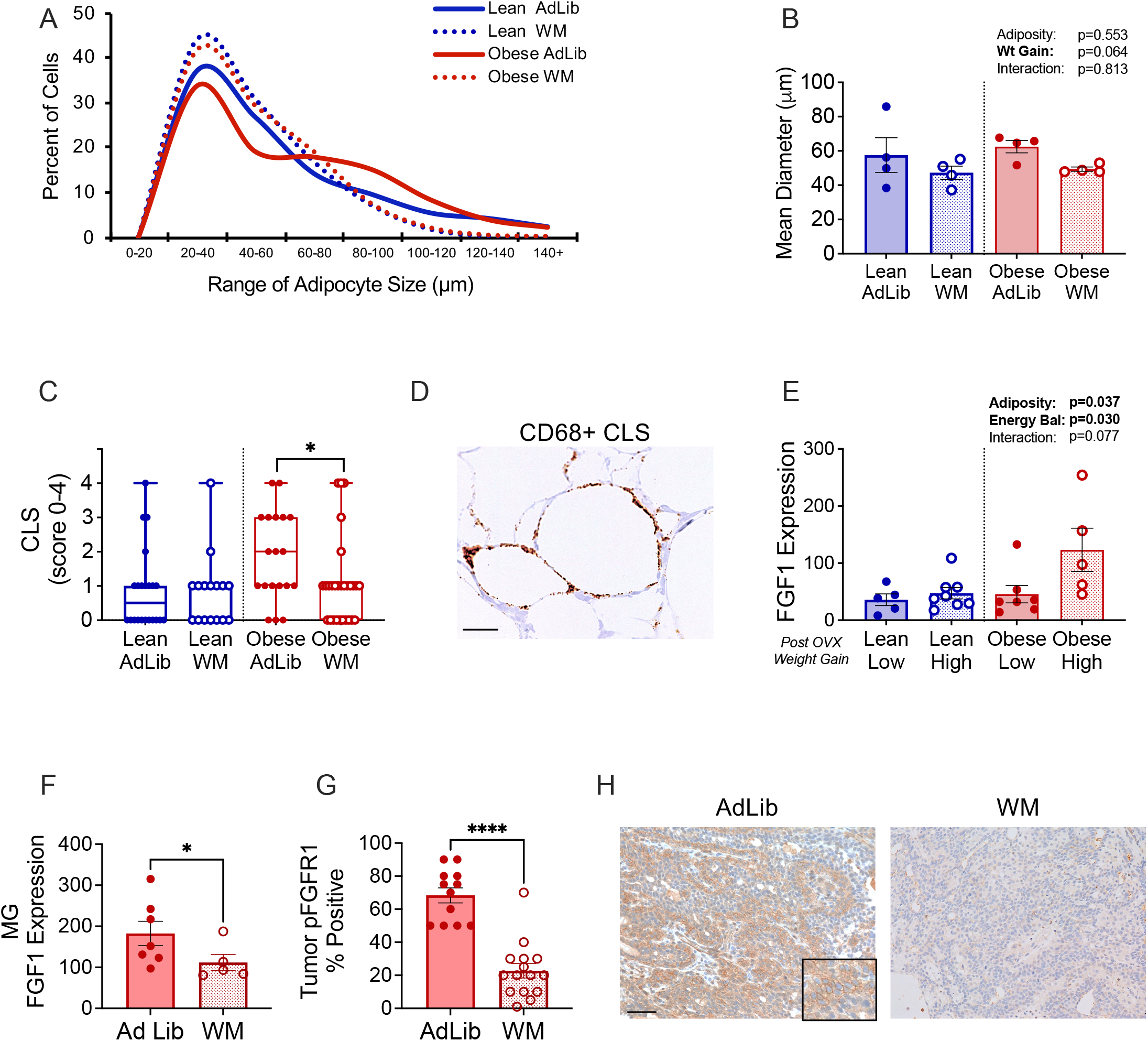
Preventing weight gain after OVX influences the tumors and microenvironment. **A)** Cell size distribution of subcutaneous/mammary adipocytes in lean and obese AdLib or WM rats at the end of study. **B)** Mean adipocyte diameter of cells in subcutaneous/mammary depots at the end of study. Two-way ANOVA, main effects of adiposity, WM, or interaction. **C)** Crown-like structure (CLS) scores, indicating local inflammation, in subcutaneous/mammary depots measured visually in blinded histological sections. **D)** Representative images of IHC staining for CD68+ CLS. **E)** Expression of Fgf1 in subcutaneous/mammary adipose tissue from lean or obese rats experiencing either a low or high rate of post-OVX weight gain and positive energy balance. Two-way ANOVA, main effects of adiposity or energy balance. **F)** Expression of Fgf1 in subcutaneous/mammary adipose tissue from obese AdLib or WM rats. Unpaired t-test, P<0.05. **G)** Levels of phosphorylated FGFR1 (pFGFR1) in the tumors of obese AdLib or WM rats evaluated by IHC and measured visually in blinded histological sections. Unpaired t-test, P<0.0001. **H)** Representative images of IHC staining for pFGFR in AdLib and WM tumors.

**Figure 5.**
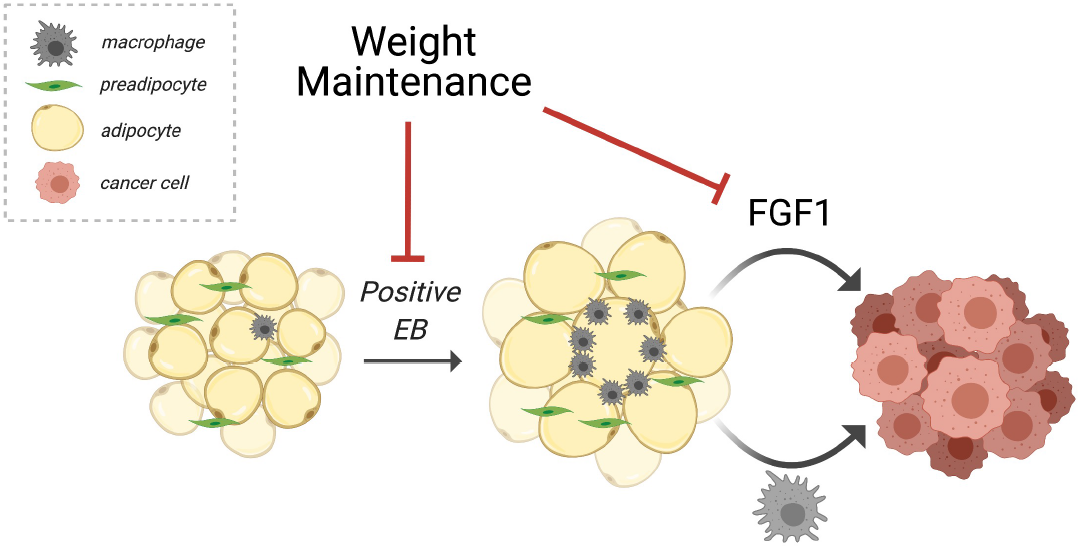
Working model diagram showing the influence of weight-gain prevention. During a positive energy balance, adipocytes become hypertrophic and rats gain weight. This associates with adipose-derived Fgf1 and inflammation in the tumor microenvironment. Both of these tumor-promotional factors are improved with weight maintenance.

To evaluate macrophage infiltration, we quantified CD68-positive crown-like structures (CLS) in mammary adipose tissue (**Fig 4C-D**). CLS are a hallmark of inflammation and dysfunctional adipose tissue that are formed as macrophages surround and engulf dying adipocytes, forming a distinct crown-like shape (38). Although the average adipocyte diameter was not significantly different between lean and obese AdLib rats (Fig 3B), there were more CLS present in mammary adipose from obese AdLib rats compared to the lean (**Fig 4C**), reflecting the shift in adipocyte size frequency distribution towards large cells. Consistent with the reduction in the number of large adipocytes in the obese WM group, CLS were also significantly reduced in mammary fat pads in obese WM rats compared to controls (**Fig 4C**). Differences were not observed in lean rats with the WM intervention, likely due to the overall lower number of CLS in this group (**Fig 4C**).

We and others have previously show that metabolic dysfunction, weight gain, and adipocyte hypertrophy are associated with increased production of FGF1 in subcutaneous adipose tissue (33,39), and activation of FGFR signaling is associated with tumor growth and resistance to endocrine therapy (33,40). Further, FGF2 has been shown to alter macrophage programming and is a critical regulator of immunity in the tumor microenvironment (41). Given the effects of weight maintenance on mammary adipocyte size and inflammation, we evaluated FGF1 levels in mammary adipose tissue from a cohort of rats in which both the adipocyte size distribution and extent of overfeeding during menopause-induced weight gain were previously measured (17). FGF1 expression was increased in obese but not lean rats experiencing a high positive energy balance (i.e. overfeeding (17)) (**Fig 4E**). FGF1 expression was lower in mammary adipose of obese WM rats compared to AdLib controls (**Fig 4F**), suggesting that the obese tumor microenvironment, in particular, has elevated inflammation and FGF1 expression after OVX that can be reduced by preventing weight gain during this time.

### Direct Effects on the Tumor

FGFs exert their growth promoting effects through binding to FGF-receptors (FGFR), which are expressed in breast cancer and immune cells. Thus, we hypothesized that higher levels of FGF in the tumor microenvironment could increase tumor growth by activating the FGF signaling pathway. We assessed FGFR phosphorylation (activation) by IHC. As shown in **Fig 4G-H**, tumors from Ob-WM rats had lower levels of phospho-FGFR1 compared to controls, consistent with lower levels of FGF1 ligand in the surrounding adipose. In addition, tumors were stained with CD68 as a marker of macrophage infiltration.

## DISCUSSION

The novel and important finding in this study is that preventing weight gain after OVX was sufficient to decrease both the growth of existing mammary tumors and the development of new tumors. This is of great significance because the menopause transition is a time when the majority of women gain weight and increase adiposity. It is notoriously difficult for women to restrict energy intake enough to lose weight during the menopause transition, as energy expenditure also decreases significantly during this window of time (42). Importantly, our data demonstrate that preventing weight gain has beneficial effects on mammary tumors, even for animals that are obese prior to OVX.

The beneficial effects of weight maintenance in this study were multi-factorial. At the systemic level, weight maintenance decreased insulin resistance and reduced visceral fat in both lean and obese rats. Circulating levels of tumor-promoting growth factors and inflammatory cytokines were also decreased, with greater effects in obese animals compared to lean. In obese rats, preventing OVX-induced weight gain also decreased adipocyte size and adipose inflammation and reduced expression of the tumor-promoting growth factor FGF-1, which associated with decreased phosphorylation of FGFR in mammary tumors, potentially contributing to the anti-tumor effects of the intervention. Taken together, these observations suggest that weight maintenance may be a viable strategy for reducing obesity-associated breast cancer risk and progression in women during the peri-/postmenopausal period.

Epidemiological data demonstrates a consistent link between obesity and postmenopausal breast cancer risk, progression, and mortality (15). Several studies have also investigated the impact of body weight changes on breast cancer risk and prognosis, beyond simply evaluating BMI. Risk estimates vary based on the population studied, but after menopause breast cancer incidence increases ~10% for every 5 BMI unit increase (i.e. transition from overweight to obese categories) (4). Adult weight gain and weight gain prior to cancer diagnosis also increase breast cancer risk and mortality, particularly for ER+ and/or progesterone receptor positive (PR+) tumors (43–47). In the European Prospective Investigation into Cancer and Nutrition (EPIC) study, long-term weight gain (>10kg) in women who were lean at age 20 was associated with a >40% increased risk of postmenopausal breast cancer compared to those who were weight stable (48). Weight gain in the year prior to, or the year after diagnosis was associated with up to a 29% increase in breast-cancer specific mortality (49). Conversely, some studies demonstrate that weight loss associates with decreased breast cancer risk. The most compelling data come from women who undergo bariatric surgery, where cancer risk is decreased by ~45% compared to untreated women, with a greater risk reduction of ER+ vs ER-tumors (50). Less extreme weight loss has also been linked to lower cancer risk; however, the exact details of these relationships vary between studies, likely due to differences in study design, duration, baseline body weight, and time during which change in body weight was assessed. For example, the WHI Observational Study (51) and the Iowa Women’s Health Study (52) both found that intentional weight loss was associated with decreased breast cancer risk, regardless of whether weight changes were monitored for a short or long time. To our knowledge, no clinical studies have yet assessed the role that menopausal weight gain plays in driving breast cancer growth. Our results showing that preventing OVX-induced weight gain had beneficial effects on all animals, regardless of their adiposity, suggest that the menopausal window may be an opportunity to lower breast cancer risk regardless of baseline adiposity at time of menopause.

While the overall effect of weight maintenance on tumor growth was the same in lean and obese rats, we found that the underlying mechanisms may vary based on adiposity. Insulin resistance was improved in both lean and obese rats, likely due in part to a reduction in visceral fat which is known to drive systemic insulin resistance. Fabian and colleagues recently reported that weight loss and visceral fat reduction improved both circulating and breast biopsy biomarkers of inflammation that have been tied to breast cancer risk (53). Women in this clinical study all had obesity at baseline; thus, it is not clear if the same benefits would be seen in non-obese women. In our preclinical study, weight maintenance had greater effects on reducing tumor-promoting growth factors and inflammatory cytokines in obese rats compared to lean. In obese rats, weight maintenance prevented mammary adipocyte hypertrophy, which has been tied to local breast inflammation and breast cancer risk (38). Our previous work in mice and human samples demonstrated that obesity and weight gain is associated with increased expression of the tumor-promoting growth factor FGF-1 (33), which is produced by hypertrophic adipocytes (39). Here, we extend these findings to show that preventing weight gain is sufficient to reduce FGF-1 in the tumor microenvironment and decrease activation of FGFR in tumors. Together, these studies suggest that weight gain leading to adipocyte hypertrophy can promote tumors through growth factor production and inflammation. It remains to be determined if targeting FGF signaling or aspects of inflammation during menopause would provide the same anticancer benefit as weight maintenance.

One limitation of our study was that surgical OVX was used as a model of menopause, which allows us to directly control the timing of the loss of ovarian hormones and manipulate body weight during this window. In women, however, menopause occurs more slowly and does not involve removal of the ovaries. We also used middle aged animals so we cannot directly assess the impact of aging in the model. Aging brings additional changes in metabolism and immune function, which may not have been identified in our middle-aged rodents. Finally, as previously mentioned, both lean and obese animals were consuming a HF diet for the duration of the study. While this removes the diet as a confounding factor in our study, it also means that the lean animals could have some level of underlying metabolic resistance greater than would be expected in many lean women.

Overall, we identified a critical life period when preventing weight gain may significantly reduce breast cancer risk and progression. Achieving sustainable weight loss is difficult for many people, including women after menopause. Our studies suggests that weight loss may not be necessary to reduce breast cancer risk and progression. Rather, preventing weight gain during a somewhat predictable life stage (menopause) may be beneficial for women and thus warrants further clinical investigation.

---

**Supplemental Table 1.**

| Timepoint | Body Weight and Composition Pre-OVX |  | p-value<sup>2</sup> |
| --- | --- | --- | --- |
|  | L<sup>1</sup> | OB<sup>1</sup> |  |
| BW 9 wks | 203.9 (5.2) | 226.1 (3.0) | <0.001 |
| BW 14 wks | 243.8 (7.0) | 277.6 (4.4) | <0.001 |
| BW OVX | 273.1 (7.2) | 353.6 (9.4) | <0.001 |
| % Fat 9 wks | 12.1 (0.5) | 14.0 (0.5) | 0.012 |
| % Fat 14 wks | 14.9 (0.6) | 19.5 (0.8) | <0.001 |
| % Fat OVX | 17.3 (0.4) | 27.6 (0.9) | <0.001 |

<sup>1</sup> Mean (std.error)<sup>2</sup> One-way ANOVA

**Supplemental Table 2:** Cumulative weight gain weeks 0 - 4 postOVX

| Characteristic | L |  | OB |  | Two-way ANOVA p-values |  |  |
| --- | --- | --- | --- | --- | --- | --- | --- |
|  | AdLib<sup>1</sup> | WM<sup>1</sup> | AdLib<sup>1</sup> | WM<sup>1</sup> | Adipo | Rx | Adipo * Rx |
| Wt gain week 0-1 | 8.8 (1.6) | -0.4 (1.6) | 7.6 (2.4) | -0.5 (2.5) | 0.6 | 0.006 | 0.8 |
| Wt gain week 0-2 | 28.0 (1.8) | 1.7 (1.4) | 24.1 (4.0) | -0.3 (2.9) | 0.3 | <0.001 | 0.8 |
| Wt gain week 0-3 | 41.8 (3.0) | 0.7 (1.5) | 38.6 (4.9) | 1.7 (2.6) | 0.5 | <0.001 | 0.6 |
| Wt gain week 0-4 | 50.9 (4.0) | 1.1 (1.2) | 50.1 (5.4) | 1.3 (2.1) | 0.9 | <0.001 | >0.9 |

<sup>1</sup> Mean (SE)

**Supplemental Table 3:** End of Study Plasma Metabolites

| Characteristic | L |  | OB |  | Two-way ANOVA p-values |  |  |
| --- | --- | --- | --- | --- | --- | --- | --- |
|  | AdLib<sup>1</sup> | WM<sup>1</sup> | AdLib<sup>1</sup> | WM<sup>1</sup> | Adipo | Rx | Adipo * Rx |
| Cholesterol, mM | 1.7 (0.1) | 1.2 (0.2) | 2.3 (0.2) | 1.8 (0.2) | 0.008 | 0.010 | >0.9 |
| Triglycerides, mM | 0.4 (0.0) | 0.4 (0.0) | 0.6 (0.0) | 0.7 (0.2) | 0.012 | 0.6 | 0.2 |
| NEFA, uM | 612.3 (43.4) | 711.7 (75.6) | 757.9 (50.3) | 763.7 (72.8) | 0.048 | 0.4 | 0.4 |

<sup>1</sup> Mean (SE)

**Supplemental Table 4:** End of Study Plasma Cytokines

| Relative Levels of Plasma Cytokines Altered with Weight Maintenance |  |  |
| --- | --- | --- |
| Cytokine | AdLib | WM |
| ACTH | 209.9 (20.6) | 109.3 (23.1) |
| Adiponectin.Acrp30 | 42,893.2 (1,818.0) | 39,426.6 (1,747.8) |
| AMPKalpha1 | 81.4 (12.2) | 22.8 (7.8) |
| BDNF | 88.9 (62.6) | 19.9 (10.6) |
| CINC2alpha.beta | 13,810.8 (1,270.4) | 21,657.6 (1,906.9) |
| CNTF | 429.0 (92.1) | 239.9 (37.3) |
| CSK | 110.6 (16.8) | 51.0 (11.5) |
| EGFR | 8.4(2.1) | 1.9 (0.5) |
| ESelectin | 721.4 (90.9) | 479.9 (53.8) |
| IL13 | 403.0 (56.6) | 134.7 (23.9) |
| IL1R6.IL1Rrp2 | 1,676.0 (662.9) | 1,351.6 (141.6) |
| IL2 | 2,384.1 (381.7) | 4,212.5 (679.5) |
| Insulin | 197.5 (43.0) | 122.2 (33.1) |
| Leptin(OB) | 210.2 (94.0) | 74.3 (21.4) |
| MIP1alpha | 1,605.2 (183.1) | 2,358.3 (235.6) |
| MIP2 | 436.8 (55.1) | 532.0 (55.0) |
| NGFR | 97.1 (14.1) | 36.8 (10.2) |
| Osteopontin.SPP1 | 70.7 (13.8) | 26.3 (7.1) |
| Thrombospondin | 373.9 (43.8) | 361.0 (141.4) |
| TIE2 | 182.9 (21.6) | 93.8 (19.9) |
| TIMP2 | 273.8 (43.8) | 119.9 (28.5) |
| TNFalpha | 144.1 (30.7) | 108.4 (38.3) |
| TRAIL | 20,125.4 (3,109.3) | 13,929.9 (3,723.6) |
| Ubiquitin | 200.1 (17.4) | 111.2 (22.9) |

## REFERENCES

1. Hales CM, Carroll MD, Fryar CD, Ogden CL. Prevalence of Obesity Among Adults and Youth: United States, 2015-2016. NCHS Data Brief 2017:1–8

2. Flegal KM, Carroll MD, Ogden CL, Curtin LR. Prevalence and trends in obesity among US adults, 1999-2008. JAMA 2010;303:235–41

3. Ogden CL, Carroll MD, Curtin LR, Lamb MM, Flegal KM. Prevalence of high body mass index in US children and adolescents, 2007-2008. JAMA 2010;303:242–9

4. Lauby-Secretan B, Scoccianti C, Loomis D, Grosse Y, Bianchini F, Straif K, et al. Body Fatness and Cancer--Viewpoint of the IARC Working Group. The New England journal of medicine 2016;375:794–8

5. Chan DS, Vieira AR, Aune D, Bandera EV, Greenwood DC, McTiernan A, et al. Body mass index and survival in women with breast cancer-systematic literature review and meta-analysis of 82 follow-up studies. Ann Oncol 2014;25:1901–14

6. Chen L, Cook LS, Tang MT, Porter PL, Hill DA, Wiggins CL, et al. Body mass index and risk of luminal, HER2-overexpressing, and triple negative breast cancer. Breast Cancer Res Treat 2016;157:545–54

7. Park YM, White AJ, Nichols HB, O’Brien KM, Weinberg CR, Sandler DP. The association between metabolic health, obesity phenotype and the risk of breast cancer. International journal of cancer 2017;140:2657–66

8. Picon-Ruiz M, Morata-Tarifa C, Valle-Goffin JJ, Friedman ER, Slingerland JM. Obesity and adverse breast cancer risk and outcome: Mechanistic insights and strategies for intervention. CA: a cancer journal for clinicians 2017;67:378–97

9. Cao Z, Zheng X, Yang H, Li S, Xu F, Yang X, et al. Association of obesity status and metabolic syndrome with site-specific cancers: a population-based cohort study. Br J Cancer 2020;123:1336–44

10. Trentham-Dietz A, Newcomb PA, Storer BE, Longnecker MP, Baron J, Greenberg ER, et al. Body size and risk of breast cancer. Am J Epidemiol 1997;145:1011–9

11. Renehan AG, Tyson M, Egger M, Heller RF, Zwahlen M. Body-mass index and incidence of cancer: a systematic review and meta-analysis of prospective observational studies. Lancet 2008;371:569–78

12. Reeves GK, Pirie K, Beral V, Green J, Spencer E, Bull D, et al. Cancer incidence and mortality in relation to body mass index in the Million Women Study: cohort study. BMJ 2007;335:1134

13. Catsburg C, Kirsh VA, Soskolne CL, Kreiger N, Bruce E, Ho T, et al. Associations between anthropometric characteristics, physical activity, and breast cancer risk in a Canadian cohort. Breast Cancer Res Treat 2014;145:545–52

14. da Silva M, Weiderpass E, Licaj I, Lissner L, Rylander C. Excess body weight, weight gain and obesity-related cancer risk in women in Norway: the Norwegian Women and Cancer study. Br J Cancer 2018;119:646–56

15. Neuhouser ML, Aragaki AK, Prentice RL, Manson JE, Chlebowski R, Carty CL, et al. Overweight, Obesity, and Postmenopausal Invasive Breast Cancer Risk: A Secondary Analysis of the Women’s Health Initiative Randomized Clinical Trials. JAMA Oncol 2015;1:611–21

16. MacLean PS, Giles ED, Johnson GC, McDaniel SM, Fleming-Elder BK, Gilman KA, et al. A surprising link between the energetics of ovariectomy-induced weight gain and mammary tumor progression in obese rats. Obesity (Silver Spring) 2010;18:696–703

17. Giles ED, Wellberg EA, Astling DP, Anderson SM, Thor AD, Jindal S, et al. Obesity and overfeeding affecting both tumor and systemic metabolism activates the progesterone receptor to contribute to postmenopausal breast cancer. Cancer Res 2012;72:6490–501

18. Giles ED, Jackman MR, MacLean PS. Modeling Diet-Induced Obesity with Obesity-Prone Rats: Implications for Studies in Females. Front Nutr 2016;3:50

19. Wellberg EA, Checkley LA, Giles ED, Johnson SJ, Oljira R, Wahdan-Alaswad R, et al. The Androgen Receptor Supports Tumor Progression After the Loss of Ovarian Function in a Preclinical Model of Obesity and Breast Cancer. Horm Cancer 2017;8:269–85

20. Giles ED, Jindal S, Wellberg EA, Schedin T, Anderson SM, Thor AD, et al. Metformin inhibits stromal aromatase expression and tumor progression in a rodent model of postmenopausal breast cancer. Breast Cancer Res 2018;20:50

21. O’Flanagan CH, Bowers LW, Hursting SD. A weighty problem: metabolic perturbations and the obesity-cancer link. Horm Mol Biol Clin Investig 2015;23:47–57

22. Young CD, Anderson SM. Sugar and fat - that’s where it’s at: metabolic changes in tumors. Breast Cancer Res 2008;10:202

23. Eliassen AH, Colditz GA, Rosner B, Willett WC, Hankinson SE. Adult weight change and risk of postmenopausal breast cancer. JAMA 2006;296:193–201

24. Radimer KL, Ballard-Barbash R, Miller JS, Fay MP, Schatzkin A, Troiano R, et al. Weight change and the risk of late-onset breast cancer in the original Framingham cohort. Nutr Cancer 2004;49:7–13

25. Pichard C, Plu-Bureau G, Neves ECM, Gompel A. Insulin resistance, obesity and breast cancer risk. Maturitas 2008

26. Jernstrom H, Barrett-Connor E. Obesity, weight change, fasting insulin, proinsulin, C-peptide, and insulin-like growth factor-1 levels in women with and without breast cancer: the Rancho Bernardo Study. Journal of women’s health & gender-based medicine 1999;8:1265–72

27. MacLean PS, Higgins JA, Giles ED, Sherk VD, Jackman MR. The role for adipose tissue in weight regain after weight loss. Obes Rev 2015;16 Suppl 1:45–54

28. Giles ED, Wellberg EA. Preclinical Models to Study Obesity and Breast Cancer in Females: Considerations, Caveats, and Tools. J Mammary Gland Biol Neoplasia 2020;25:237–53

29. Thompson HJ, McGinley JN, Wolfe P, Singh M, Steele VE, Kelloff GJ. Temporal sequence of mammary intraductal proliferations, ductal carcinomas in situ and adenocarcinomas induced by 1-methyl-1-nitrosourea in rats. Carcinogenesis 1998;19:2181–5

30. MacLean PS, Higgins JA, Johnson GC, Fleming-Elder BK, Peters JC, Hill JO. Metabolic adjustments with the development, treatment, and recurrence of obesity in obesity-prone rats. Am J Physiol Regul Integr Comp Physiol 2004;287:R288–97

31. Giles ED, Jackman MR, Johnson GC, Schedin PJ, Houser JL, MacLean PS. Effect of the estrous cycle and surgical ovariectomy on energy balance, fuel utilization, and physical activity in lean and obese female rats. Am J Physiol Regul Integr Comp Physiol 2010;299:R1634–42

32. Young S, Hallowes RC. Tumours of the mammary gland. IARC Sci Publ 1973:31–73

33. Wellberg EA, Kabos P, Gillen AE, Jacobsen BM, Brechbuhl HM, Johnson SJ, et al. FGFR1 underlies obesity-associated progression of estrogen receptor-positive breast cancer after estrogen deprivation. JCI Insight 2018;3

34. Kabos P, Finlay-Schultz J, Li C, Kline E, Finlayson C, Wisell J, et al. Patient-derived luminal breast cancer xenografts retain hormone receptor heterogeneity and help define unique estrogen-dependent gene signatures. Breast Cancer Res Treat 2012;135:415–32

35. Greendale GA, Han W, Finkelstein JS, Burnett-Bowie SM, Huang M, Martin D, et al. Changes in Regional Fat Distribution and Anthropometric Measures Across the Menopause Transition. J Clin Endocrinol Metab 2021;106:2520–34

36. Neeland IJ, Ayers CR, Rohatgi AK, Turer AT, Berry JD, Das SR, et al. Associations of visceral and abdominal subcutaneous adipose tissue with markers of cardiac and metabolic risk in obese adults. Obesity (Silver Spring) 2013;21:E439–47

37. Fang L, Guo F, Zhou L, Stahl R, Grams J. The cell size and distribution of adipocytes from subcutaneous and visceral fat is associated with type 2 diabetes mellitus in humans. Adipocyte 2015;4:273–9

38. Iyengar NM, Gucalp A, Dannenberg AJ, Hudis CA. Obesity and Cancer Mechanisms: Tumor Microenvironment and Inflammation. J Clin Oncol 2016;34:4270–6

39. Wang S, Cao S, Arhatte M, Li D, Shi Y, Kurz S, et al. Adipocyte Piezo1 mediates obesogenic adipogenesis through the FGF1/FGFR1 signaling pathway in mice. Nat Commun 2020;11:2303

40. Formisano L, Stauffer KM, Young CD, Bhola NE, Guerrero-Zotano AL, Jansen VM, et al. Association of FGFR1 with ERalpha Maintains Ligand-Independent ER Transcription and Mediates Resistance to Estrogen Deprivation in ER(+) Breast Cancer. Clin Cancer Res 2017;23:6138–50

41. Im JH, Buzzelli JN, Jones K, Franchini F, Gordon-Weeks A, Markelc B, et al. FGF2 alters macrophage polarization, tumour immunity and growth and can be targeted during radiotherapy. Nat Commun 2020;11:4064

42. Lovejoy JC, Champagne CM, de Jonge L, Xie H, Smith SR. Increased visceral fat and decreased energy expenditure during the menopausal transition. Int J Obes (Lond) 2008;32:949–58

43. van den Brandt PA, Ziegler RG, Wang M, Hou T, Li R, Adami HO, et al. Body size and weight change over adulthood and risk of breast cancer by menopausal and hormone receptor status: a pooled analysis of 20 prospective cohort studies. Eur J Epidemiol 2021;36:37–55

44. Azrad M, Blair CK, Rock CL, Sedjo RL, Wolin KY, Demark-Wahnefried W. Adult weight gain accelerates the onset of breast cancer. Breast Cancer Res Treat 2019;176:649–56

45. Vrieling A, Buck K, Kaaks R, Chang-Claude J. Adult weight gain in relation to breast cancer risk by estrogen and progesterone receptor status: a meta-analysis. Breast Cancer Res Treat 2010;123:641–9

46. Cleveland RJ, Eng SM, Abrahamson PE, Britton JA, Teitelbaum SL, Neugut AI, et al. Weight gain prior to diagnosis and survival from breast cancer. Cancer Epidemiol Biomarkers Prev 2007;16:1803–11

47. Chan DSM, Abar L, Cariolou M, Nanu N, Greenwood DC, Bandera EV, et al. World Cancer Research Fund International: Continuous Update Project-systematic literature review and meta-analysis of observational cohort studies on physical activity, sedentary behavior, adiposity, and weight change and breast cancer risk. Cancer Causes Control 2019;30:1183–200

48. Ellingjord-Dale M, Christakoudi S, Weiderpass E, Panico S, Dossus L, Olsen A, et al. Longterm weight change and risk of breast cancer in the European Prospective Investigation into Cancer and Nutrition (EPIC) study. Int J Epidemiol 2021

49. Chan DSM, Vieira AR, Aune D, Bandera EV, Greenwood DC, McTiernan A, et al. Body mass index and survival in women with breast cancer-systematic literature review and meta-analysis of 82 follow-up studies. Ann Oncol 2014;25:1901–14

50. Feigelson HS, Caan B, Weinmann S, Leonard AC, Powers JD, Yenumula PR, et al. Bariatric Surgery is Associated With Reduced Risk of Breast Cancer in Both Premenopausal and Postmenopausal Women. Ann Surg 2020;272:1053–9

51. Chlebowski RT, Luo J, Anderson GL, Barrington W, Reding K, Simon MS, et al. Weight loss and breast cancer incidence in postmenopausal women. Cancer 2019;125:205–12

52. Parker ED, Folsom AR. Intentional weight loss and incidence of obesity-related cancers: the Iowa Women’s Health Study. Int J Obes Relat Metab Disord 2003;27:1447–52

53. Fabian CJ, Klemp JR, Marchello NJ, Vidoni ED, Sullivan DK, Nydegger JL, et al. Rapid Escalation of High-Volume Exercise during Caloric Restriction; Change in Visceral Adipose Tissue and Adipocytokines in Obese Sedentary Breast Cancer Survivors. Cancers 2021;13:4871

